# Detecting Epistasis with the Marginal Epistasis Test in Genetic Mapping Studies of Quantitative Traits

**DOI:** 10.1101/066985

**Authors:** Lorin Crawford, Ping Zeng, Sayan Mukherjee, Xiang Zhou

## Abstract

Epistasis, commonly defined as the interaction between multiple genes, is an important genetic component underlying phenotypic variation. Many statistical methods have been developed to model and identify epistatic interactions between genetic variants. However, because of the large combinatorial search space of interactions, most epistasis mapping methods face enormous computational challenges and often suffer from low statistical power due to multiple test correction. Here, we present a novel, alternative strategy for mapping epistasis: instead of directly identifying individual pairwise or higher-order interactions, we focus on mapping variants that have non-zero *marginal epistatic effects* — the combined pairwise interaction effects between a given variant and all other variants. By testing marginal epistatic effects, we can identify candidate variants that are involved in epistasis without the need to identify the exact partners with which the variants interact, thus potentially alleviating much of the statistical and computational burden associated with standard epistatic mapping procedures. Our method is based on a variance component model, and relies on a recently developed variance component estimation method for efficient parameter inference and p-value computation. We refer to our method as the “MArginal ePIstasis Test”, or MAPIT. With simulations, we show how MAPIT can be used to estimate and test marginal epistatic effects, produce calibrated test statistics under the null, and facilitate the detection of pairwise epistatic interactions. We further illustrate the benefits of MAPIT in a QTL mapping study by analyzing the gene expression data of over 400 individuals from the GEUVADIS consortium.

**Author Summary:** Epistasis is an important genetic component that underlies phenotypic variation and is also a key mechanism that accounts for missing heritability. Identifying epistatic interactions in genetic association studies can help us better understand the genetic architecture of complex traits and diseases. However, the ability to identify epistatic interactions in practice faces important statistical and computational challenges. Standard statistical methods scan through all-pairs (or all high-orders) of interactions, and the large number of interaction combinations results in slow computation time and low statistical power. We propose an alternative mapping strategy and a new variance component method for identifying epistasis. Our method examines one variant at a time, and estimates and tests its *marginal epistatic effect* — the combined pairwise interaction effects between a given variant and all other variants. By testing for marginal epistatic effects, we can identify variants that are involved in epistasis without the need of explicitly searching for interactions. Our method also relies on a recently developed variance component estimation method for efficient and robust parameter inference, and accurate p-value computation. We illustrate the benefits of our method using simulations and real data applications.

## Introduction

Genetic mapping studies, in the form of genome-wide association studies (GWASs) [1] and molecular trait quantitative trait loci (QTL) mapping studies [2–5], have identified thousands of genetic loci associated with many complex traits and common diseases, providing insights into the genetic basis of phenotypic variation. Most of these existing genetic mapping studies look at one variant at a time and focus on identifying marginal genetic associations that exhibit either additive or dominant effects. However, it has long been hypothesized that effects beyond additivity could contribute to a large proportion of phenotypic variation. In particular, epistasis — the interaction between genetic loci — is thought to play a key role in defining the genetic architecture underlying complex traits [6, 7] and constituting the genetic basis of evolution [8, 9]. Indeed, studies have detected pervasive epistasis in many model organisms [10–33]. However, substantial controversies remain [34–36]. For example, in some settings, genetic mapping studies have identified many candidates of epistatic interactions that contribute to quantitative traits and diseases [37–40], but some of these effects can be explained by additive effects of other unsequenced variants [41]. On the other hand, while previous variance partition studies have shown that genetic variance for many traits are mainly additive [34, 35, 42], these conclusions have been challenged recently [36]. Furthermore, while modeling epistasis has been recently shown to increase phenotype prediction accuracy in modal organisms [43] and facilitate genomic selection in animal breeding programs [44, 45], such conclusions do not hold in all settings [46]. Finally, epistasis has been recently proposed as one of the main factors that explain missing heritability — the proportion of heritability not explained by the top associated variants in GWASs [1, 47]. In particular, studies have hypothesized that epistasis can confound heritability estimation in pedigree studies and cause inflation of heritability estimates, creating the so-called “phantom heritability” [48,49]. However, for some traits, the contribution of epistasis to missing heritability is negligible [50].

Nevertheless, because of the potential importance of epistasis in defining the genetic architecture of complex traits, many statistical methods have been developed to identify epistatic interactions in genetic mapping studies [51, 52]. Different existing statistical methods differ in their ways of selecting a testing unit (i.e. variants or genes [53]), their searching strategy (e.g. exhaustive search [54–56] or probabilistic search [57] or prioritization based on a candidate set [58]), and the calculation of test statistics (e.g. various frequentist tests [59] or Bayesian approaches [60–62]). However, almost all of these statistical methods focus on explicitly searching for pairwise or higher-order interactions when identifying epistatic effects. Because of the extremely large search space (e.g. *p*(*p* − 1)/2 pairwise combinations for *p* variants), these methods often suffer from heavy computational burden and low statistical power. Despite various efficient computational implementations [56, 63] and recently developed efficient search algorithms [57], exploring over a large combinatorial search space remains a daunting task for large epistasis mapping studies. Statistically, because of a lack of *a priori* knowledge of epistatic loci, exploring all combinations of genetic variants could result in low statistical power — on the other hand, restricting to a subset of prioritized combinations based on prior knowledge or marginal effects could also miss important genetic interactions.

Here, we present an alternative strategy for mapping epistasis. Instead of directly identifying individual pairwise or higher-order interactions, we focus on identifying variants that have a non-zero interaction effect with any other variants. To do so, we develop a novel statistical method, which we refer to as the the “MArginal ePIstasis Test” (MAPIT), to test each variant in turn on its *marginal epistatic effect* — the combined pairwise interaction effects between a given variant and all other variants. By testing marginal epistatic effects, we can identify candidate markers that are involved in epistasis without the need to identify the exact partners with which the variants interact — thus, potentially alleviating much of the statistical and computational burden associated with standard epistatic mapping methods. In addition, evidence of marginal epistasis can be used to further prioritize the search and identification of pairwise interactions. Our method is based on variance component models [64–76]. By taking advantage of a recently developed variance component estimation method [77] for efficient parameter inference and p-value computation, our method is scalable to moderately sized genetic mapping studies. We illustrate how MAPIT can serve as a useful alternative to standard methods in mapping epistasis with both simulations and a real data application.

## Methods and Material

### MAPIT Model

We describe the MArginal ePIstasis Test in detail here. Our goal is to identify variants that interact with other variants, and to avoid explicitly searching for pairwise interactions. Therefore, unlike standard tests for epistasis, MAPIT works by examining one variant at a time. For the *k*^th^ variant, we consider the following linear model,

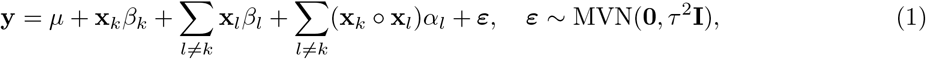

where **y** is an *n*-vector of phenotypes for *n* individuals; *µ* is an intercept term; **x***_k_* is an *n*-dimensional genotype vector for the *k*^th^ variant that is the focus of the model; *β_k_* is the corresponding additive effect size; **x***_l_* is an *n*-dimensional genotype vector for the *l*^th^ variant, and *l* represents any of the *p* variants other than the *k*^th^; *β_l_* is the corresponding additive effect size; **x***_k_* ○ **x***_l_* denotes an element-wise multiplication between genotype vectors, thus representing the interaction term between the *k*^th^ and *l*^th^ variants; *α_l_* is the corresponding interaction effect size; ***ε*** is an *n*-vector of residual errors; *τ* ^2^ is the residual error variance; **I** is the identity matrix; and MVN denotes a multivariate normal distribution. In addition, we assume that the genotype vector for each variant has been centered and standardized to have mean 0 and standard deviation 1.

The model in Equation (1) is an underdetermined linear system (*p > n*). Therefore, we have to make additional modeling assumptions on the effect sizes *β_l_* and *α_l_* to make the model identifiable. To do so, we follow standard approaches [64, 67, 69] and assume that each individual effect size follows a normal distribution, or 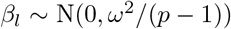 and 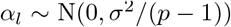 for *l* ≠ *k*. With the normal assumption on effect sizes, the model in Equation (1) is equivalent to the following variance component model,

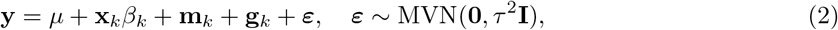

where 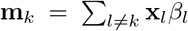 is the combined additive effects from all other variants, and effectively represents the additive effect of the *k*^th^ variant under the polygenic background of all other variants; 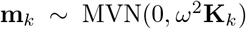 with 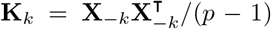 being the genetic relatedness matrix computed using genotypes from all variants other than the 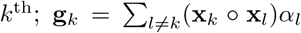 is the summation of all pairwise interaction effects between the *k*^th^ variant and all other variants; and **g***_k_* ~ MVN(0*, σ*^2^**G***_k_*) with **G***_k_* = **D***_k_***K***_k_***D***_k_* representing a relatedness matrix computed based on pairwise interaction terms between the *k*^th^ variant and all other variants. Here, we denote **D***_k_* = diag(**x***_k_*) to be an *n × n* diagonal matrix with the genotype vector **x***_k_* as its diagonal elements. It is important to note that both **K***_k_* and **G***_k_* change with every new marker *k* that is considered.

We want to point out that the formulation of MAPIT in Equation (2) can also be easily extended to accommodate other fixed effects (e.g. age, sex, or genotype principal components), as well as other random effects terms that can be used to account for sample non-independence due to other genetic or common environmental factors. In addition, we choose to model *β_k_* as a fixed effect here, but modeling it as a random effect is straightforward. Also note that, in this work, we limit ourselves to only consider second order epistatic relationships between SNPs. However, the generalization of MAPIT to detect higher order interactions is straightforward and only involves the manipulation of **G***_k_*.

### Point Estimates

Our goal is to identify variants that have non-zero interaction effects with any other variant. To do so, we can examine each variant in turn (*k* = 1,…, *p*) and test the null hypothesis in Equation (1) that variant *k* has no interaction effect with any other variant, 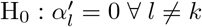. This same null hypothesis is specified in the variance component model stated in Equation (2) as H_0_ : *σ*^2^ = 0. The variance component *σ*^2^ effectively captures the total epistatic interaction effects between the *k*^th^ variant and all other variants — we call this the marginal epistatic effect for the *k*^th^ variant.

Testing the marginal epistatic effect *σ*^2^ requires jointly estimating the variance component parameters (*ω*^2^, *σ*^2^, *τ* ^2^) in Equation (2). The standard method for variance component estimation is the restricted maximum likelihood estimation (REML) method. However, REML is computationally slow: it requires an iterative optimization procedure where the time complexity of each iteration scales cubically with the number of individuals [66,68–73]. The slow computation speed of REML is further exacerbated by the fact that the variance component model changes for every variant *k* (i.e. both **K***_k_* and **G***_k_* are variant specific) — hence, the variance component parameters are required to be estimated over and over again across genome-wide variants. Therefore, we cannot use REML for marginal epistatic mapping. Instead, we follow the recently developed MQS method [77] for efficient variance component estimation and testing. MQS is based on the method of moments and produces estimates that are mathematically identical to the Haseman-Elston (HE) cross-product regression [78]. Note that MQS is not only computationally more efficient than HE regression, but also provides a simple, analytic estimation form that allows for exact p-value computation — thus alleviating the need for jackknife re-sampling procedures [79] that both are computationally expensive and rely on incorrect individual independence assumptions [80].

To estimate the variance components with MQS, we first multiply a projection matrix **M***_k_* on both sides of the model in Equation (2) to remove the influence of *µ* and **x***_k_*. Here, 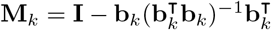, where **b***_k_* = [**1**_*n*_, x_*k*_] with **1***_n_* denoting an *n*-vector of ones. Thus, **M***_k_* is a variant specific projection matrix onto both the null space of the intercept and the corresponding genotypic vector **x***_k_*. By multiplying **M***_k_*, we obtain the following simplified modeling specification

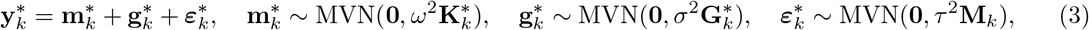

where 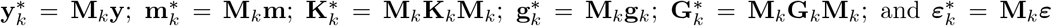, respectively. Note that Equation (3) also changes with every new marker *k* that is considered.

To simplify notation, we use ***δ*** = (*ω*^2^*, σ*^2^*, τ* ^2^) to denote the variance components. Next, we use the notation 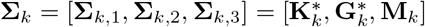. Lastly, we use indices *i, j, l* ∈ {1, 2, 3} to represent the corresponding variance component or covariance matrix. Given estimates 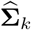, we can obtain the MQS estimates for the variance components of each variant 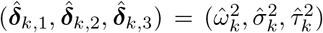 via the following simple analytic formula

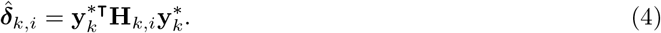

Here, we define 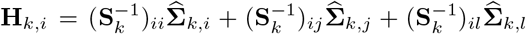, where **S***_k_* is a 3 × 3 matrix in which 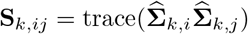 for every *i, j, l* = 1, 2, 3.

### Hypothesis Testing

MAPIT provides two options to compute p-values. The first option is approximate, and is based on a normal test that only requires the variance component estimate 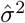 and its corresponding standard error. In particular, the variances of the MQS estimates in Equation (4) are given via a previously suggested and computationally efficient approximation [77]

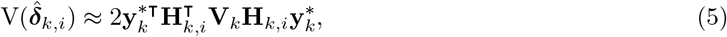

where 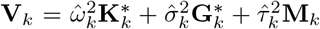. Given an estimate from Equation (4) and its standard error from Equation (5), we can perform a normal test (or z-test) to compute p-values. More specifically, we use a two sided test since the MQS estimates can be either positive or negative. The normal test is computationally efficient, but it is important to stress that when the sample size is small it is not appropriate.

We also provide a second, exact option to compute p-values which is valid in the cases of small sample sizes. This second option relies on the fact that the MQS variance component estimate in Equation (4) follows a mixture of chi-square distributions under the null hypothesis. This is because **y**\* is assumed to follow a multivariate normal distribution under the modeling assumptions. In particular, 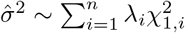, where 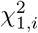 are chi-square random variables with one degree of freedom and (*λ*_1_*, …, λ_n_*) are the corresponding eigenvalues of the matrix

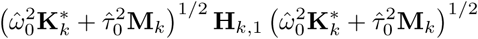

with (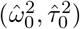 being the MQS estimates of (*ω*^2^, *τ* ^2^) under the null hypothesis. We can then use the Davies method [64, 81] to compute p-values.

While the Davies method is the appropriate test of choice and is expected to produce calibrated p-values, it can become computationally demanding as the numbers of observed samples becomes large (see S1 Table). Specifically, the computational complexity of the normal test scales linearly with the number of markers and quadratically with the number of individuals. On the other hand, the computational complexity of the Davies method scales linearly with the number of markers, but cubically with the number of individuals. Therefore, the Davies method can be much slower than the normal test. For example, while analyzing 10,000 markers on a data set with 1,000, 2,500 and 5,000 individuals, the z-test version of MAPIT requires an average of 2.1, 18.9, and 67.6 minutes, respectively. Using the Davies method on these same sets of data, MAPIT requires about 6.1, 108.6, and 654.9 minutes, respectively. Therefore, in practice, we advertise a hybrid p-value computation procedure that uses the normal test by default, and then applies the Davies method when the p-value from the normal test is below the threshold of 0.05. The hybrid procedure combines the advantages of the two different tests and produces calibrated p-values while remaining computationally efficient (again see S1 Table). As we will also show in the results section, the above MQS estimation and testing procedures allow for both accurate and efficient marginal epistatic mapping in moderately sized genetic mapping studies.

### Other Methods

In this work, we compare MAPIT to two different epistatic mapping approaches. The first is a single-SNP additive association analyses which is fit with a linear regression model by using the `-lm` argument in the GEMMA software [66–68]. This software is publicly available at http://www.xzlab.org/software.html. The second identifies pairwise interactions directly by implementing an exhaustive search linear model, which we fit by using the `--epistasis` argument in the PLINK software (version 1.9) [82]. This software is also publicly available at https://www.cog-genomics.org/plink2/epistasis.

Besides estimating and testing marginal epistatic effects for every SNP, we also explore the use of a variance component model to estimate the total contribution of pairwise epistasis onto the phenotypic variance [44, 83, 84]. More specifically, we consider a linear mixed model which partitions the total phenotypic variance using variance components corresponding to additive and pairwise epistatic covariance matrices:

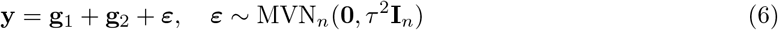

where **g**_1_ ~ MVN*_n_*(**0**, *ω*^2^**K**) is the linear effects component; **g**_2_ ~ MVN*_n_*(**0**, *σ*^2^**K**^2^) is the pairwise interaction component; and ***ε*** represents the proportion of phenotypic variance explained by random noise. Here, we let {*ω*^2^, *σ*^2^, *τ*^2^} be corresponding random effect variance terms. The matrix **I***_n_* is an *n × n* identity matrix. The covariance matrix **K** = **XX**^T^/*p* is the conventional (linear) genetic relatedness matrix, as previous defined. The covariance matrix **K**^2^ = **K** ○ **K** represents a pairwise interaction relationship matrix, and is obtained by using the Hadamard product (i.e. the squaring of each element) of the linear kernel matrix with itself [44, 83, 84].

In the presence of population structure and potential stratification effects, we modify Equation (6) by including the top 10 genotype principal components as fixed effects. The modified variance component model can be fitted through the following steps. First, we collect top 10 PCs in a covariate matrix **Z**. Next, we compute a projection matrix **M** = **I − Z**(**Z**^T^**Z**)^−1^**Z**^T^, and multiply it on both sides of the model in Equation (6) to get the following transformed model:

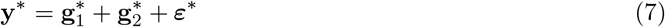

where 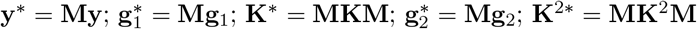; and 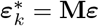, respectively.

We apply the first procedure to simulations, and the second procedure to both the simulations and real data analyses. In either case, our goal is to estimate the influence of the total contribution across pairwise epistatic effects have on the phenotype. We quantify these contributions by examining the proportion of phenotypic variance explained (pPVE) using the following equation defined in [66, 77] for every variance component *i*:

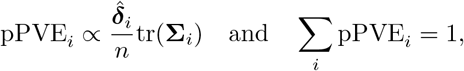

where **Σ** = [**K**\*, **K**^2^*, **M**]. In the current study, with Equation (7), *i* = 1, …, 3. We specifically calculate the pPVEs corresponding to the random effect variance terms 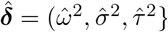. The variance component that explains the greatest proportion of the overall PVE then represents the most influential effect onto that particular phenotypic response. We implement this model with multiple variance components in the current study by using the `-vc` argument within the GEMMA software. We also use the standard output (i.e. point estimate and standard error) to conduct an asymptotic normal test to assess the power of this approach.

### Software Availability

The software implementing MAPIT is freely available at https://github.com/lorinanthony/MAPIT. We use the CompQuadForm R package to compute p-values from the Davies method. The Davies method can sometimes yield a p-value equal exactly to 0 when the true p-value is extremely small [85]. In this case, we report p-values as *P* ≈ 0. If this is of concern, one can compute the p-values for MAPIT using Kuonen's saddlepoint method [85, 86] or Satterthwaite's approximation equation [87].

### Real Data Sets

In the present study, we utilize two real data sets: one from the Wellcome Trust Case Control Consortium (WTCCC), and the other from the GEUVADIS Consortium Project. For the first, we specifically used the control samples from the WTCCC 1 study [88] (http://www.wtccc.org.uk/), which consists of 2,938 individuals with 458,868 SNPs following the quality control steps described in detail in a previous study [67].

For the second, we obtained the GEUVADIS data [4] (http://www.geuvadis.org) which contains gene expression measurements for 462 individuals from five different populations: CEPH (CEU), Finns (FIN), British (GBR), Toscani (TSI) and Yoruba (YRI). Following previous studies [89], we focused only on protein coding genes and lincRNAs that are annotated from GENCODE (release 12) [90]. We removed lowly expressed genes that had zero counts in at least half of the individuals, and obtained a final set of 15,607 genes. Afterwards, following previous studies [89], we performed PEER normalization [91] to remove confounding effects and unwanted variations. In order to remove potential population stratification, we quantile normalized the gene expression measurements across individuals in each population to a standard normal distribution, and then quantile normalized the gene expression measurements to a standard normal distribution across individuals from all five populations. In addition to the gene expression data, all individuals in GEUVADIS also have their genotypes sequenced in the 1000 Genomes project [92]. Among the sequenced genotypes, we retained 1,236,922 SNPs that have a minor allele frequency (MAF) above 0.05 and missingness below 0.01. Then, for each gene in turn, we obtained its *cis*-SNPs that are located within either 100 kb upstream of the transcription start site (TSS) or 100 kb downstream of the transcription end site (TES), resulting in a total of 2,735,891 unique SNP-gene combinations with an average of 175 *cis*-SNPs per gene.

In the GEUVADIS data set, we perform four sets of analyses. The first involves using MAPIT with a genetic relatedness matrix **K***_cis_*, where for the expression of each gene **K***_cis_* was computed using only the corresponding *cis*-SNPs. The second, involves using MAPIT with a genetic relatedness matrix **K***_trans_*, where for the expression of each gene **K***_trans_* is computed using only corresponding *trans*-SNPs located outside of the defined 100 kb *cis*-window. The third analysis corresponds to using MAPIT with a genome-wide genetic relatedness matrix **K***_GW_*, where **K***_GW_* was computed using all SNPs in the study. Besides these analyses, we also performed a fourth analysis to guard against potential residual population stratification. For this analysis, we first compute the top 10 principal components from the genotype matrix, and then collect them in a matrix **Z**. Next, we regress this matrix of confounding factors onto the expression of each gene, and save the residuals — with which we perform another quantile normalization. We also save the residuals of the genotype matrix after removing the effects of the confounding factors and compute a relatedness matrix **K***_Pop_* based on the genotype residuals. Finally, we implement MAPIT on the normalized expression residuals using **K***_Pop_*.

## Results

### Simulations: Type I Error Control

To validate MAPIT and our proposed hybrid testing procedure, in terms of controlling type I error, we carried out a simulation study. Specifically, we utilize the genotypes from chromosome 22 of the control samples in the WTCCC 1 study [88] to generate continuous phenotypes. Exclusively considering this group of individuals and SNPs leaves us with an initial dataset consisting of *n* = 2,938 control samples and *p* = 5,747 markers.

In order to investigate the type I error control, we first subsample from the genotypes for *n* = 1,000, 1,750, and 2,500 subjects. Next, we randomly select 1,000 causal SNPs and simulate continuous phenotypes by using the following two simulation models: (i) a standard model with **y** = **X*β*** + ***ε***, and (ii) a population stratification model with **y** = **Zu** + **X*β*** + ***ε***, where **X** is the genotype matrix, **Z** contains covariates representing population structure, and **u** are fixed effects. Under the first model, we simulate both the additive effect sizes of each causal SNP and the random noise term from a standard normal distribution, and then we scale the two terms further to ensure a narrow-sense heritability of 60%. In the second model, we introduce population stratification effects into the simulations by allowing the top 5 and 10 genotype principal components (PCs) to make up 10% of the overall phenotypic variance (i.e. through the **Zu** term). These population stratification effect sizes are also drawn from a standard normal distribution. Note that, for both settings, the idea of the null model holds because there are no interaction effects, and MAPIT solely searches for significant marginal epistatic effects that are a summation of pairwise interactions. Furthermore, in the cases in which simulations were conducted under model (ii), the genotype PCs were not included while running MAPIT, and no other preprocessing normalization procedures were carried out to account for the added population structure. All evaluations of calibration and type 1 error are strictly based on the linear mixed model presented in Equation (2).

We assess the calibration of MAPIT under both the normal test and the Davies method for each sample size *n*. Figure 1 shows the quantile-quantile (QQ) plots based on simulation model (i), with the application of MAPIT to these null datasets under both hypothesis testing strategies. Similar QQ-plots for data simulated under model (ii) can be found in Supporting Information (see S1 Fig.). The normal test heavily relies on the assumption of asymptotic normality — therefore, it is expected to see improvement of performance as the sample size increases. However, as one also expects, the normal test is inaccurate in the extreme tails of the test even for larger sample sizes — hence, the inflation of the normal test p-values in Figure 1. Alternatively, utilizing the Davies method via a mixture of chi-squares allows MAPIT to robustly control for type I error across all sample sizes — even in the presence of population stratification effects. MAPIT's ability to produce calibrated type I error in the presence of population stratification is not surprising, as the model of MAPIT contains a genetic relatedness matrix that has been well known to effectively control for population stratification [66, 70, 76]. Table 1 shows the empirical type I error rates estimated for MAPIT at significance levels *α* = 0.05, 0.01, and 0.001, respectively, for simulations under model (i). Again, similar tables for data simulated under mode (ii) can be found in Supporting Information (see S2 and S3 Table). As expected based on the QQ-plots under the Davies method, MAPIT controls the type I error rate for reasonably sized datasets, and can be slightly liberal when the sample size is small. Presumably, the liberal behavior of p-values in small samples arises from the fact that frequentist tests do not account for uncertainty in the variance component estimates in the null model. Based on the null simulation results, the Davies method should be the choice of default. However, for computational reasons, we use a hybrid p-value computation procedure (details in Methods and Material) that recalibrates p-value for a SNP using the Davies method when the z-test p-value for the SNP is below the nominal threshold of 0.05. The results we present throughout the rest of the paper will be based on using MAPIT with this hybrid approach.

**Figure 1.**
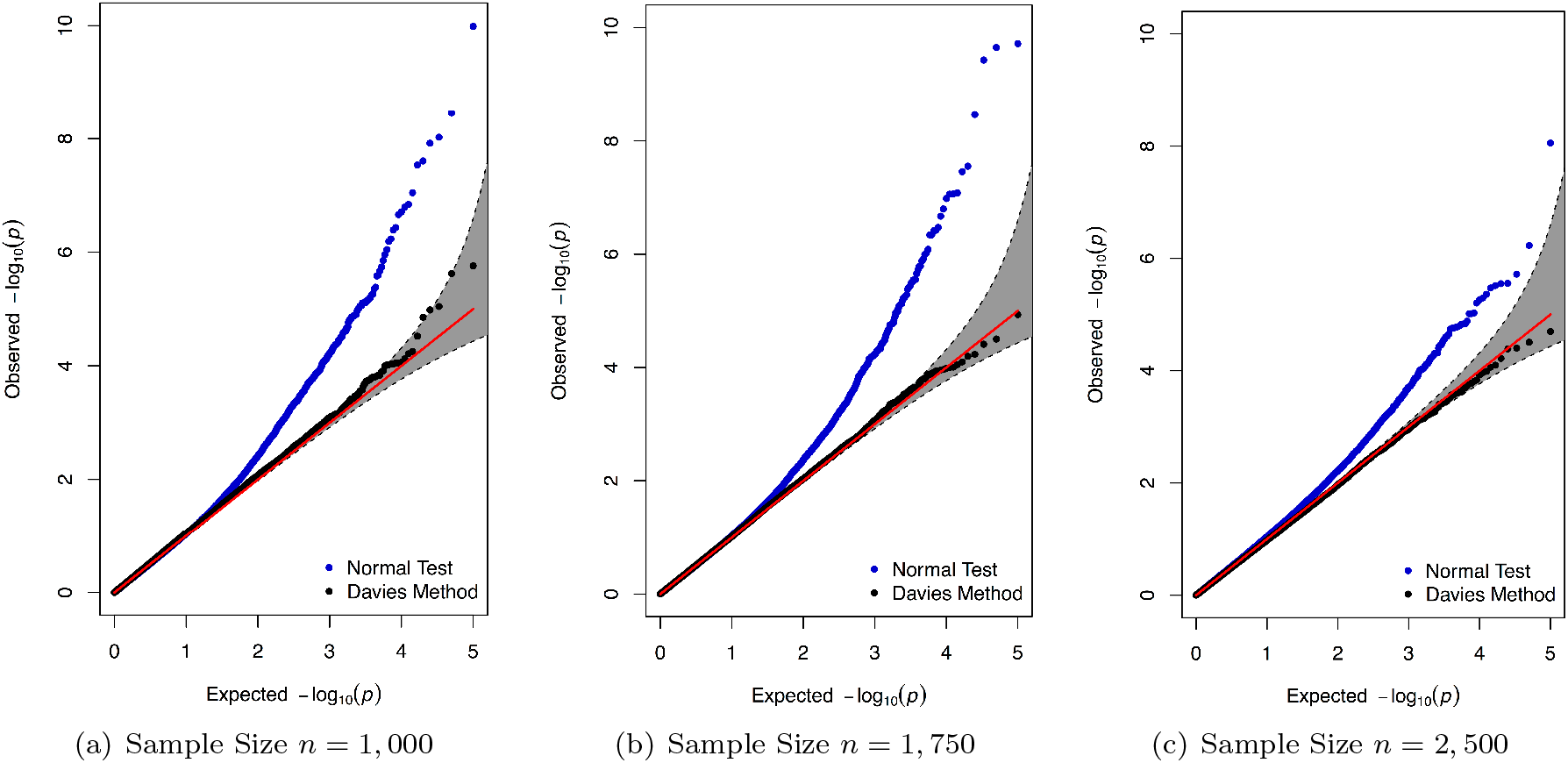
Calibration of p-values produced by MAPIT via QQ-plots. The QQ-plots applying MAPIT to 100 simulated null datasets assuming sample sizes (a) 1,000, (b) 1,750, and (c) 2,500. Blue dots are p-values produced by under the normal test (or z-test), while the black dots represent p-values tested using the Davies method via a mixture of chi-square distributions. The 95% confidence intervals for the null hypothesis of no association are shown in grey.

**Table 1.**
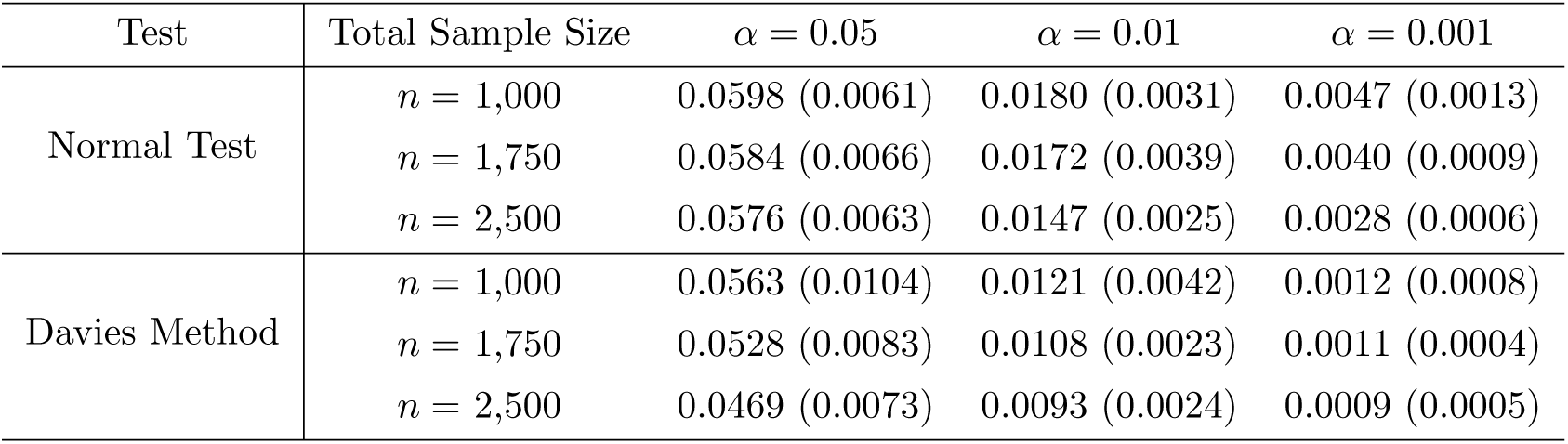
Empirical type I error estimates of MAPIT. Each entry represents type I error rate estimates as the proportion of p-values a under the null hypothesis based on 100 simulated continuous phenotypes for the normal test (or z-test) and the Davies method. These results are based on 100 simulated data sets using simulation model (i). Empirical size for the analyses used significance thresholds of *α* = 0.05, 0.01, and 0.001. Sample sizes were set to 1,000, 1,750, and 2,500. Values in the parentheses are the standard deviations of the estimates.

### Simulations: Estimating and Identifying Marginal Epistatic Effects

In this section, we use simulation studies to illustrate the advantages of MAPIT in identifying marginal epistatic associations. In addition, besides correctly detecting marginal epistatic associations, we will show that MAPIT can also estimate the marginal epistatic effects reasonably well. Therefore, analogous to SNP heritability estimation settings [67, 69, 75], these variance component estimates can serve as a measurement of the marginal interaction phenotypic variance explained (PVE) by each epistatic causal variant [77].

To test the power of MAPIT, we again consider simulation designs similar to those proposed by previous epistatic analysis studies [63]. First, we assume that the broad-sense heritability is known (H^2^ = 0.6) [67, 88, 93]. Next, we use the 22^nd^ chromosome of all control cases from the WTCCC 1 study **X** (i.e. *n* ≈ 3,000 and *p* ≈ 6,000) to simulate continuous phenotypes that mirror genetic architectures affected by a combination of additive and pairwise epistatic effects. Specifically, we randomly choose 1,000 causal SNPs to directly affect the phenotype and classify the causal variants into three groups: (1) a small set of interaction SNPs, (2) a larger set of interaction SNPs, and (3) a large set of additive SNPs. In the simulations carried out in this study, SNPs interact between sets, so that SNPs in the first group interact with SNPs in the second group, but do not interact with variants in their own group (the same rule applies to the second group). One may view the SNPs in the first set as the “hubs” in an interaction map. We are reminded that interaction (epistatic) effects are different from additive effects. All causal SNPs in both the first and second groups have additive effects and are involved in pairwise interactions, while causal SNPs in the third set only have additive effects.

The additive effect sizes of all causal SNPs again come from a standard normal distribution or ***β*** ~ MVN(**0**, **I**). Next, we create a separate matrix **W** which holds the pairwise interactions of all the causal SNPs between groups 1 and 2. These SNPs have effect sizes also drawn as ***α*** ~ MVN(**0**, **I**). We scale both the additive and pairwise genetic effects so that collectively they explain a fixed proportion of genetic variance. Namely, the additive effects make up *ρ*%, while the pairwise interactions make up the remaining (1 − *ρ*)%. Once we obtain the final effect sizes for all causal SNPs, we draw errors to achieve the target H^2^. The phenotypes are then created by summing all effects using two simulation models: (i) **y** = **X*β*** + **W*α*** + ***ε*** and (ii) **y** = **Zu** + **X*β*** + **W*α*** + ***ε***, where **Zu** again represents population stratification. In the latter model, population stratification effects are introduced into the simulations by allowing the top 5 and 10 genotype principal components (PCs) **Z** to make up 10% of the overall variation in the trait. The effect sizes for these stratification effects are also drawn as **u** ~ MVN(**0**, **I**).

We consider a few scenarios that depend on two parameters:

- (1 − *ρ*), which measures the portion of H^2^ that is contributed by the interaction effects of the first and second groups of causal SNPs. Specifically, the phenotypic variance explained (PVE) by the additive genetic effects is said to be V(**X*β***) = *ρ*H^2^, while the PVE of the pairwise epistatic genetic effects is given as V(**W*α***) = (1 − *ρ*)H^2^.
- *p*_1_/*p*_2_/*p*_3_, which are the number of causal SNPs in each of the three groups, respectively.

Specifically, we set *ρ* = {0.5, 0.8} and choose *p*_1_/*p*_2_/*p*_3_ = 10/10/980 (scenario I), 10/20/970 (scenario II), 10/50/940 (scenario III), and 10/100/890 (scenario IV). Note that scenarios III and IV assume a larger number of interactions than scenario I and II do, and are thus likely to be closer to reality. The particular case where *ρ* = 0.5, the additive and epistatic effects are assumed to equally contribute to the broad-sense heritability of the simulated phenotypes. The alternative case in which *ρ* = 0.8 is a case where the PVE of the simulated complex traits are dominated by additive effects. We analyze 100 different simulated datasets for each value of *ρ*, across each of the four scenarios. All of the results described in this section are based on the cases in which phenotypes were simulated under model (i) with *ρ* = 0.8, as this case is a more realistic setting for human traits where epistatic effects only make up a small percentage of the broad-sense heritability. The results for *ρ* = 0.5 can be found in Supporting Information (see S2 Fig.). Similar results for all data simulated under model (ii) can also can be found in Supporting Information (see S3 and S4 Fig.) Once again, note that in the cases for which simulations were conducted under model (ii), the genotype PCs were not included while running MAPIT, and no other preprocessing normalization procedures were carried out to account for the added population structure. All evaluations of MAPIT are strictly based on the linear mixed model presented in Equation (2).

Figure 2(a) shows the power results for MAPIT's ability to detect both group 1 and 2 causal variants, respectively, compared across each simulation scenario. Empirical power of MAPIT was estimated as the proportion of p-values below 0.05. We can see MAPIT's ability to detect both groups of causal markers depends on the pairwise interaction PVE explained by each variant. For example in Figure 2(a), each causal variant in group 1 is expected to explain V(**W*α***)/*p*_1_ = 1.2% of the true interaction PVE since in every scenario *p*_1_ = 10. In these situations, the cumulative PVE of these markers is great and MAPIT's power is large for all four scenarios (approximately 30% power). Note that this power is similar to MAPIT's ability to detect the group 2 causal markers under Scenario I (i.e. *p*_2_ = 10), where each epistatic variant is also expected to explain V(**W*α***)/*p*_2_ = 1.2% of the interaction PVE. Alternatively, MAPIT exhibits half of the power when detecting the group 2 SNPs in the case of Scenario II (i.e. *p*_2_ = 20), as each SNP explains only V(**W*α***)/*p*_2_ = 0.6% of the PVE (approximately 15% power). In addition, MAPIT's power to identify group 1 variants is independent of the number of variants in group 2 (i.e. *p*_2_), suggesting that MAPIT's power depends on the total interaction effects, rather than individual pairwise effects or the number of interacting pairs. The results based on the genome-wide significance threshold are similar and can be found in Supporting Information (see S5 and S6 Fig.).

**Figure 2.**
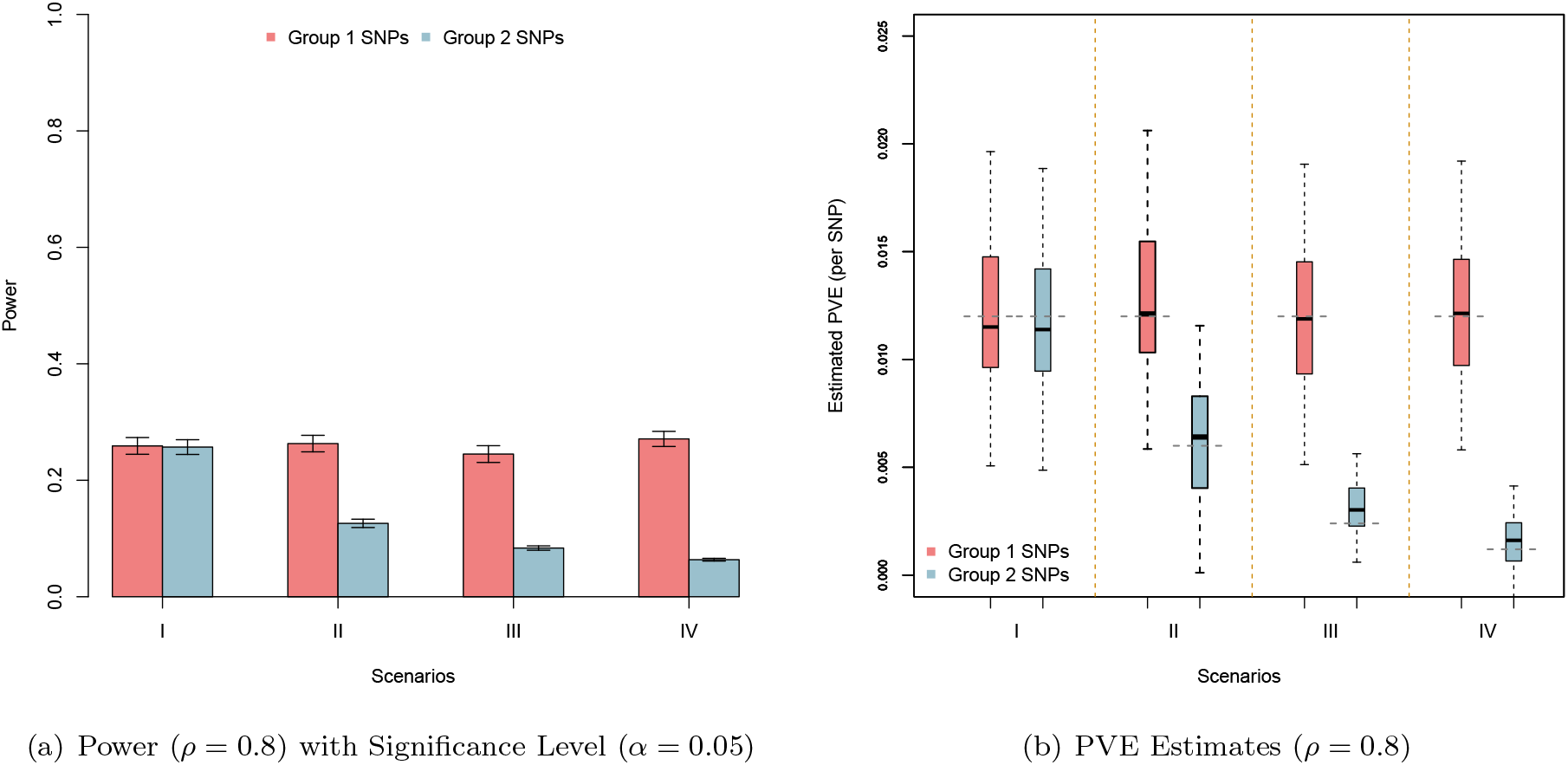
Empirical power to detect simulated causal interacting makers and estimating their marginal PVE. Groups 1 and 2 causal markers are colored in light red and light blue, respectively. These figures are based on a broad-sense heritability level of H^2^ = 0.6 and parameter *ρ* = 0.8, estimated with 100 replicates. Here, *ρ* = 0.8 was used to determine the portion of broad-sense heritability contributed by interaction effects. Figure (a) shows the power of MAPIT to identify SNPs in each causal group under significance level *α* = 0.05. The lines represent 95% variability due to resampling error. Figure (b) shows boxplots of the marginal PVE estimates for the group 1 and 2 causal SNPs from MAPIT for the four simulation scenarios. The true PVEs per causal SNP (0.012 for the group 1 SNPs; 0.012, 0.006, 0.0024, and 0.0012 for the Group 2 SNPs) are shown as dashed grey horizontal lines.

While our main focus is on testing and identifying epistasis, we also assess MAPIT's ability to estimate the contribution of each group 1 and 2 causal SNP to the interaction PVE. Figure 2(b) show boxplots of these estimates. The true interaction PVE explained by each causal SNP is depicted as the grey dashed lines. These plots show that even though MAPIT's power is directly affected by the epistatic contribution to the phenotypic variation, its ability to correctly estimate the effects of causal interacting SNPs is robust and approximately unbiased. It is important to note that we see MAPIT maintain its estimation ability even when the portion of PVE explained by a set of causal SNPs is very small (i.e. group 2 SNPs in scenario IV). The estimation results are consistent with the well-known robustness of variance component models in estimating PVE in other settings (e.g. estimation of SNP heritability) [67, 69, 75]. Finally, further deviating from our main focus, we also apply a standard variance component model to partition the phenotypic variance into an additive component and an epistatic component following the approach of [44, 83, 84] (details in Methods and Material). Results show that the standard variance component model can also be used to estimate the total contribution of epistasis reasonably well (see S7 Fig.), and produces reasonable power at the significance level of 0.05 with a standard asymptotic normal test (see S8 Fig.).

### Simulations: Power Comparisons

Here, we compare the performance of MAPIT with a standard exhaustive search procedure that examines all pairwise interactions to explicitly identify the exact pairs of variants involved in epistatic interactions [55,56]. Specifically, for the exhaustive search, we consider the PLINK linear model **y** = *µ* + **x***_i_β_i_* + **x***_j_β_j_* + (**x***_i_*◦**x***_j_*)*α_ij_* +***ε*** and test H_0_ : *α_ij_* = 0 for every marker combination of *i* and *j* in turn [82]. Keeping notation consistent, **x***_i_* ◦ **x***_j_* denotes element-wise multiplication between genotypes *i* and *j*, and *α_ij_* represents the effect size of their interaction. Note that the exhaustive search procedure is computationally feasible within PLINK because we only have *p* ≈ 6, 000 markers in the simulations.

It is helpful to point out here that the purpose of this comparison is to depict MAPIT as a viable alternative for the exhaustive search procedure. We will show that MAPIT not only can perform a significance test to detect variants involved in epistasis, but also can be used to obtain a prioritized set of variants that are further used to identify pairwise interactions. Our simulation comparisons are thus targeted to illustrate how MAPIT can be used in these two tasks, and how its performance differs from the exhaustive search procedure in different scenarios.

#### Identifying variants involved in epistasis

We first compare MAPIT against the PLINK exhaustive search method in identifying variants that are involved in epistasis. For this task, MAPIT can directly perform a significance test and produce a p-value. Here, we note that the power of MAPIT and the exhaustive search method are determined by different factors: the power of the linear interaction method depends on each individual epistatic interaction effect size *α_ij_*, while the power of MAPIT, as we have shown in the previous section, depends on the marginal epistatic effects — the summation of interaction effects. Therefore, we would expect MAPIT and the exhaustive search method to be advantageous in different situations (if the exhaustive search method is computationally feasible). In particular, we would expect the exhaustive search method to be more powerful in scenario I (and II) where each individual interaction effect is large, and MAPIT to be more powerful in scenario (III and) IV where each individual interaction effect is small but the marginal epistatic effect remains large. Furthermore, MAPIT can naturally account for population stratification via the included genetic relatedness matrix, while the exhaustive search method requires including genotype PCs as covariates to control for such confounding effects. Therefore, we would also expect MAPIT to be robustly more powerful when there are confounding population stratification effects that are not explicitly taken into account. To validate our expectations, we again generate continuous outcomes using the same two previously described simulation schemes: (i) **y** = **X*β*** + **W*α*** + ***ε*** and (ii) **y** = **Zu** + **X*β*** + **W*α*** + ***ε***. Once again, all results described in the main text are based on model (i) with *ρ* = 0.8, while all other results can be found in Supporting Information (see S9-S13 Fig.).

We evaluate MAPIT's and PLINK's ability to accurately identify marginal epistatic effects for markers in each of the two causal groups. The criteria we use compares the false positive rate (FPR) with the rate at which true variants are identified for each model (TPR). Figure 3 depicts the ability of MAPIT and PLINK to detect causal variants in groups 1 and 2. In particular, these plots depict the portion of causal markers discovered after prioritizing all of those considered in order of their significance. We assess the marginal epistatic detection in the PLINK exhaustive search by first running the previously described pairwise linear model, ordering the resulting p-values for each possible interaction, and drawing a power curve for identifying the SNPs that are members of simulated causal groups 1 and 2. For example, if the top p-values from the exhaustive search are interactions SNP1-SNP2, SNP2-SNP3, SNP4-SNP5, and only SNP2 is the true causal epistatic variant, then the top three pairs only marginally identify 1 true variant and 4 false variants.

**Figure 3.**
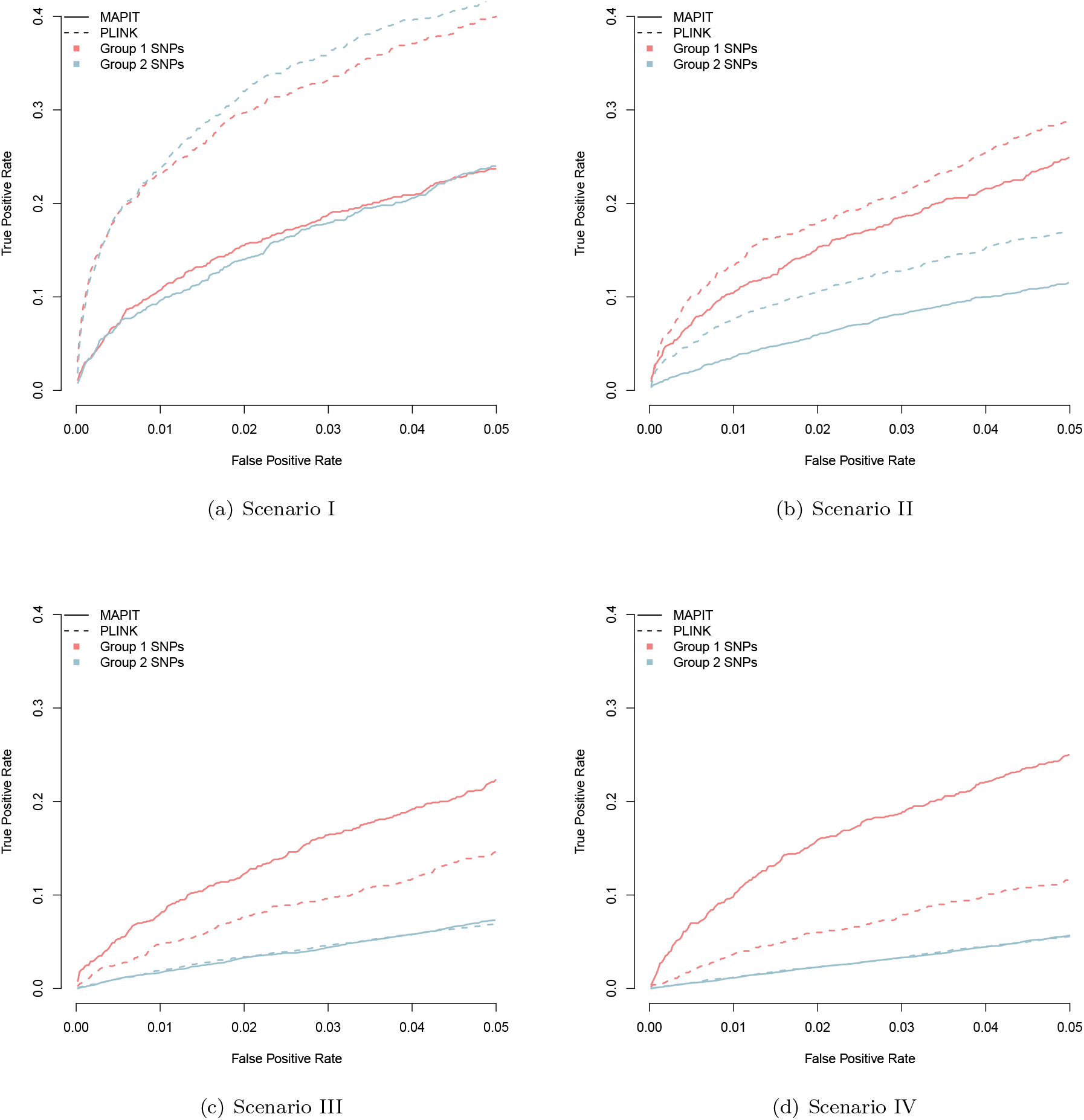
Power analysis for detecting group 1 and group 2 causal SNPs. We compare the mapping abilities of MAPIT (solid line) to the exhaustive search procedure in PLINK (dotted line) in all scenarios (alternating panels), under broad-sense heritability level H^2^ = 0.6 and *ρ* = 0.8. Here, *ρ* = 0.8 was used to determine the portion of broad-sense heritability contributed by interaction effects. Group 1 (light red) and group 2 (light blue) causal SNPs. The x-axis shows the false positive rate, while the y-axis gives the rate at which true causal variants were identified. Results are based on 100 replicates in each case.

As expected, while the power of MAPIT depends on the pairwise interaction PVE explained by each SNP, the power of the exhaustive search depends on the individual interaction effect size. For example, the power of the exhaustive search to detect group 1 causal epistatic SNPs is dependent on the number of group 2 causal SNPs, which also determines the interaction effect size in simulations. Therefore, while the exhaustive search exhibits higher power in the sparse scenario where there are only a small number of interactions each with a large effect size (e.g. scenarios I and II), its power quickly decays in the more polygenic scenario where there is a large number of interactions, each with a small effect size (e.g. scenarios III and IV). MAPIT is able to perform well in the more realistic polygenic scenarios (III and IV) by modeling the marginal epistatic effects of each variant, allowing the detection of epistatic variants not to be dependent on the individual pairwise interaction effect size. Importantly, MAPIT remains powerful even in the presence of population stratification, as it can effectively control for population stratification with the included genetic relatedness matrix. In contrast, the exhaustive search approach in PLINK does not explicitly control for population stratification, and can suffer from power loss in the presence of stratification especially when the total epistatic contribution (i.e. 1 − *ρ*) is small (see again S11 and S13 Fig.).

We are reminded that another advantage to MAPIT is the reduced space it must search over, as MAPIT only requires *p* tests for a data set with *p* genetic markers. Therefore, MAPIT is expected to exhibit a computational advantage over this type of exhaustive search approach in moderate size genetic mapping studies with millions of markers.

#### Identifying pairwise interactions

Although we have focused on identifying marginal epistatic effects so far, MAPIT can also be used to facilitate the identification of pairwise (or high-order) epistatic interactions. In addition to comparing MAPIT with the exhaustive search method from PLINK, we also consider another common approach for identifying epistatic pairs — a two-step filtering association mapping procedure [51, 55, 58]. These types of filtering methods often first apply a marginal (additive) single-SNP test to identify associated genetic variants with non-zero additive effects, and then focus on the identified variants to test all pairwise interactions between them. Depending on the correlation between the marginal additive effect size and the probability of being involved in epistasis with SNPs genome-wide, these filtering methods can be more powerful than the exhaustive search strategy mentioned in the previous section — not to mention they are certainly much more computationally efficient. However, for any trait, because there is no expectation that SNPs involved in epistasis will always have large additive effects, these filtering methods will not always outperform the exhaustive search method, and can sometimes be significantly under powered (as our simulations and real data application will show). Here, instead of using the additive test, we propose using MAPIT as the initial filter. We hypothesize that the initial list of associated SNPs from MAPIT will be more robust and more likely capture epistatic effects, as MAPIT directly prioritizes SNPs based on marginal epistatic effects. By using marginal epistatic evidence in the initial filtering step, we expect MAPIT to outperform the previous common procedure of using a linear model for filtering.

In this set of simulations, we utilize the same subset of real genotypes used for the marginal epistatic simulations in the last section [88], and again generate phenotypes under the same four simulation scenarios with the same two simulation models where pairwise interactions are well defined. After randomly selecting the three sets of causal SNPs and creating their pairwise interactions, we run MAPIT and a single-SNP additive linear model (via GEMMA) [66] using all variants. We also reuse the PLINK exhaustive search, again as a baseline comparison. For MAPIT and the single-SNP linear model in GEMMA, we rank each variant according to their marginal p-values. The top 100 SNPs identified by both models are then selected, and all pairwise interactions among them are tested using a linear model that controls for the two main effects. For the PLINK exhaustive search, we simply rank the top 100^2^ interactions to assess pairwise power.

Figure 4 compares the power of the filtering procedures using the two different methods as an initial step. Phenotypes used to create this figure were generated under each scenario with broad-sense heritability H^2^ = 0.6. As in previous sections, all results described in this section are based on model (i) with *ρ* = 0.8, and all other results can be found in Supporting Information (see S14 and S15 Fig.). Compared with the single-SNP test, filtering SNPs using MAPIT provides more power in finding true pairwise epistatic interactions. In fact, even for the cases in which the marginal additive effects contribute to a majority of the broad-sense heritability (i.e. *ρ* = 0.8), using MAPIT as the initial filtration procedure (as opposed to the single-SNP additive linear model) provides more power for finding exact causal epistatic pairs. This improvement comes from the fact that MAPIT allows the ranking of variants to be based on their marginal epistatic effects, rather than their marginal additive effects. Therefore, the set of SNPs identified by MAPIT in the first step already contains variants that capture epistatic effects, thus resulting in higher power in the second step to identify epistatic interaction pairs. In addition, similar to the simulation comparison in the previous subsection, MAPIT and the exhaustive search procedure are advantageous in different settings: the exhaustive search procedure is again more powerful in the sparse setting where each individual pairwise interaction is large (i.e. scenarios I and II) while MAPIT gains an advantage in the polygenic setting where there a large number of interactions each with small effects (i.e. scenarios III and IV). Again, in the presence of population stratification, MAPIT remains powerful while the exhaustive search procedure suffers from substantial power loss (see again S15 Fig.). Overall, MAPIT also represents an attractive alternative to identifying pairwise interactions.

**Figure 4.**
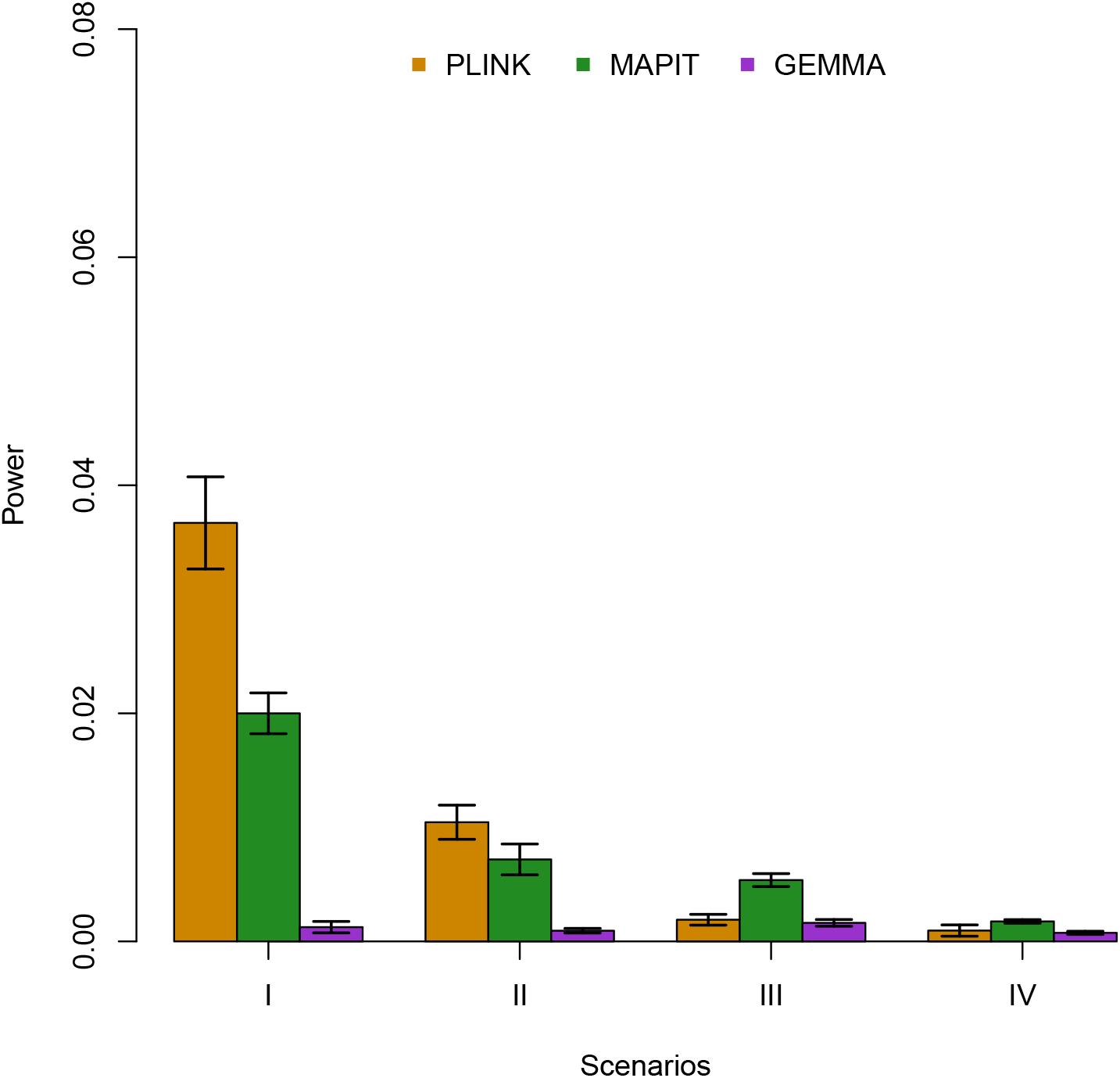
Empirical power of exhaustive search procedures to detect epistatic pairs. Here, the effectiveness of MAPIT (green) as an initial step in a pairwise detection filtration process is compared against the more conventional single-SNP testing procedure, which is carried out via GEMMA (purple). In both cases, the search for epistatic pairs occurs between the top 100 significant marginally associated SNPs are considered. We use the fully exhaustive search model in PLINK (orange) as a baseline comparison. We compare the three methods in all scenarios (x-axis), under broad-sense heritability level H^2^ = 0.6. Here, *ρ* = 0.8 was used to determine the portion of broad-sense heritability contributed by interaction effects. The y-axis gives the rate at which true causal epistatic pairs were identified. Results are based on 100 replicates in each case. The lines represent 95% variability due to resampling error.

### Detecting Epistasis in GEUVADIS

We assess MAPIT's ability to detect epistasis in a quantitative trait loci (QTL) association mapping study for gene expression levels (i.e. eQTL study). Often times, eQTL studies deal with SNP effect sizes (on gene expression levels) that are orders of magnitude larger than that (on organism-level traits) from GWASs [2–5], thus facilitating in the identification of epistasis. Indeed, recent studies have started to reveal an initial set of epistatic interactions that underlie gene expression variation [37, 94, 95]. By applying MAPIT to eQTL studies, we hope to better understand the genetic architecture that underlie gene expression variation.

The specific data set that we consider in this section features 462 individuals from five different populations whose gene expression data was collected by the GEUVADIS consortium [4]. These individuals also have their genotypes sequenced by the 1000 Genomes project [92]. In order to identify potential QTLs involved in associated pairwise epistatic interactions, we exclusively use MAPIT to analyze variants that passed quality control filters and were located within 100 kb of each gene of interest (Materials and Methods). Such variants represent likely *cis*-acting QTL, which are more readily identifiable in small sample sizes than trans-QTL [2–5]. Overall, we apply MAPIT to a final data set that consists of approximately 16,000 genes, 1.2 million SNPs, and 2.7 million SNP-gene combinations.

To remove population stratification and other confounding effects, we have removed top factors from the gene expression matrix and normalized the gene expression data within each of the five populations separately — all before performing a final joint normalization [4] (details in Methods and Material). The results of MAPIT presented in this section are based on using additive and epistatic relatedness matrices derived from a covariance matrix **K***_cis_*, where for the expression of each gene **K***_cis_* was computed using only the corresponding *cis*-SNPs. While using **K***_cis_*, MAPIT tests the summed epistatic effects between a given *cis*-SNP and all other *cis*-SNPs within the same gene. The primary reason that we present the main results based on **K***_cis_* is to allow for a fair comparison with the exhaustive search method (details below) — which, due to computational reasons, can only be applied to examine all *cis*-pairs. To assess whether or not these initial results are sensitive to the choice of covariance matrix, we also implement MAPIT using a genetic relatedness matrix **K***_trans_*, where for the expression of each gene **K***_trans_* was computed using only the corresponding *trans*-SNPs located outside the *cis*-window (Methods and Material). Using MAPIT with **K***_trans_* assesses the summed epistatic effects between a given *cis*-SNP and all other *trans*-SNPs. Lastly, we consider the implementation of MAPIT with a genome-wide genetic relatedness matrix **K***_GW_*, where **K***_GW_* was computed using all SNPs in the study (Methods and Material). Here, using MAPIT with **K***_GW_* tests the summed epistatic effects between a given *cis*-SNP and all SNPs genome-wide. Intuitively, using **K***_cis_* will be more powerful than using either **K***_trans_* or **K***_GW_* if epistatic interactions are more likely to happen between *cis*-SNP pairs, rather than between *cis*-SNP and genome-wide SNP pairs — however, less powerful otherwise. In addition to these three sets of analyses, to guard against any potential residual population stratification, we also removed the effects of the top 10 genotype principal components and used the residual expression data together with a **K***_Pop_* computed using the genome-wide residual genotype data for a further analysis. The purpose of the additional analyses is to highlight the robustness of the results found using MAPIT.

To contrast MAPIT's marginal association findings, we also directly compare results from the single-SNP additive model via GEMMA and the fully exhaustive search model in PLINK. From this point forward, we will refer to QTL identified by MAPIT as marginally epistatic QTL (*mepiQTL*), the QTL detected by GEMMA as the more conventional expression QTL (*eQTL*), and the QTL found by PLINK as epistatic QTL (*epiQTL*). Similarly, we will refer to genes that have at least one mepiQTL as *mepiGenes*, genes that have at least one eQTL as *eGenes*, and genes that have at least one epiQTL as *epiGenes*.

In this analysis, a significant marginal association for a particular SNP identified by MAPIT or GEMMA was determined by using a gene specific Bonferroni-corrected significance p-value threshold *P* = 0.05/∑*s_i_* = 1.828 × 10^−8^, where *s_i_* is the number of *cis*-SNPs for gene *i*. For PLINK, a single SNP was deemed marginally significant if it belonged to a epistatic pair with a p-value below the genome-wide threshold *P* = 1.09 × 10^−10^ (i.e. Bonferroni correction for a total of 455,801,241 examined cis-SNP pairs). While using the genetic relatedness matrix **K***_cis_*, MAPIT identified a total of 3,434 mepiQTL across 228 different mepiGenes that satisfied this marginal significance rule. Additionally, while using the covariance matrix **K***_trans_*, MAPIT detected a total of 2,160 across 130 mepiGenes. Similarly, MAPIT also identified a total of 2,160 mepiQTL across 130 mepiGenes when using **K***_GW_* (i.e. identical to that identified by **K***_trans_*), and this number changed to 3,056 mepiQTL across 184 different mepiGenes after correcting for residual population stratification and using **K***_Pop_*. GEMMA, on the other hand, found 55,645 significant eQTL across 1,417 different eGenes, and no significant eQTL or eGenes after correction for potential residual population stratification. Note that the former number of eGenes as detected by GEMMA is similar to that from the original research [4], with slight differences most likely due to data set variations (e.g. data origins, preprocessing measures, etc.). Lastly, PLINK identified 14,722 significant epiQTL spanning across 99 epiGenes prior to accounting for any residual confounding effects, and 17,286 significant epiQTL spanning across 102 epiGenes after the extra correction. The amount of overlap between the mepiQTL/mepiGenes detected by MAPIT, the eQTL/eGenes identified by GEMMA, and the epiQTL/epiGenes found by PLINK is explicitly specified in Supporting Information (see S4 and S5 Tables).

The histogram of the MAPIT p-values for all SNP-gene combinations, while using **K***_cis_*, is presented in Figure 5(a) with a corresponding QQ-plot of the log_10_ p-values presented in Supporting Information (see S16(a) Fig.). Both of these figures illustrate strong signals on a background of uniformly distributed p-values. Similar results for MAPIT with **K***_trans_*, **K***_GW_*, and **K***_Pop_* can be found in Supporting Information (see S16(b), S16(c), S16(d), S17(a), S17(b), and S17(c) Fig.). S6-S9 Tables also list the p-values for all significant mepiQTL as computed via a MAPIT scan over the *cis*-windows of each gene, with and without correction for population stratification. The distribution of locations for mepiQTL and eQTL, relative to the gene transcription start site (TSS) and the gene transcription end site (TES), is depicted in Figure 6 (see also S18 Fig.). Consistent with evaluations of other QTL association mapping studies [2–5], eQTLs detected by GEMMA are mostly enriched near TSS. In contrast, mepiQTLs are enriched near the TES in addition to the TSS, with distance to genes showing a slightly wider spread pattern than eQTLs.

**Figure 5.**
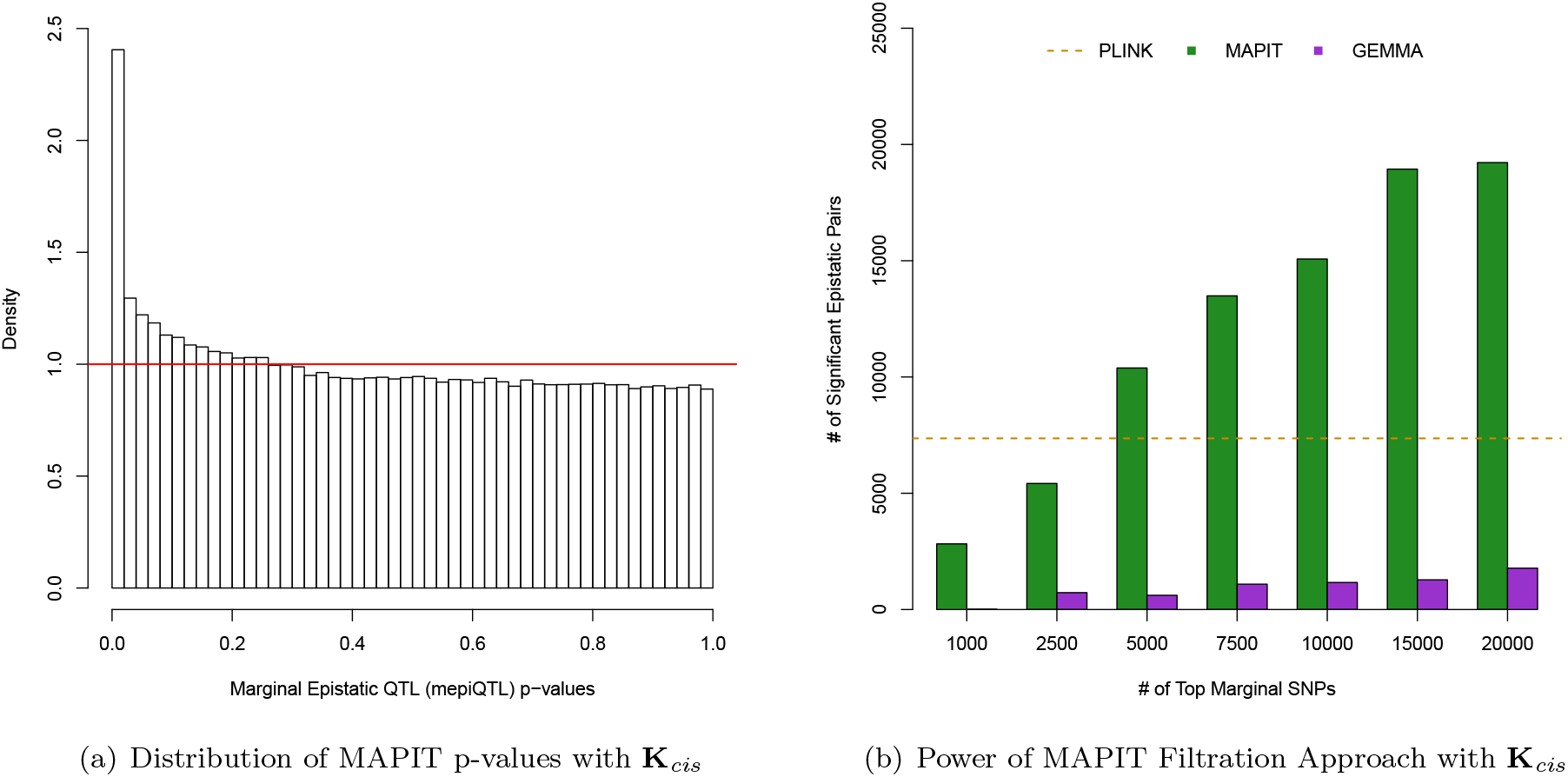
Comparison of epistatic filtration methods with MAPIT and GEMMA on the GEUVADIS data set. All of these results are based on using MAPIT with genetic relatedness matrix **K***_cis_*. Figure (a) shows a histogram of the MAPIT p-values for all variants in the GEUVADIS data set. The horizontal red line corresponds to a uniform distribution of p-values. Figure (b) shows the number of significant pairwise interactions (y-axis) identified by MAPIT (green) and GEMMA (purple) when searching between the top {1000, 2500, 5000, 7500, 10000, 15000, 20000} marginally associated variants (x-axis). We use the number of significant pairs identified by fully exhaustive search model in PLINK as a baseline comparison (orange dotted line). This image shows the distributions of genome-wide significant epistatic pairs as found by each method. An interaction for MAPIT and GEMMA was deemed signifiant if it had a joint p-value below the threshold *P* = 0.05/(∑*_i_ q_i_*(*q_i_* − 1)/2), where *q_i_* is the number of top variants located in the *cis*-window of gene *i*. In the case of PLINK, we consider two variants to be a significantly associated epistatic pair if they have a joint p-value below the threshold *P* = 1.09 × 10^−10^, which corresponds to the Bonferroni-correction that would be used if we examined all possible genome-wide SNP pairs across all genes in the final data set. Overall, PLINK detected 7,361 significant epistatic pairs.

**Figure 6.**
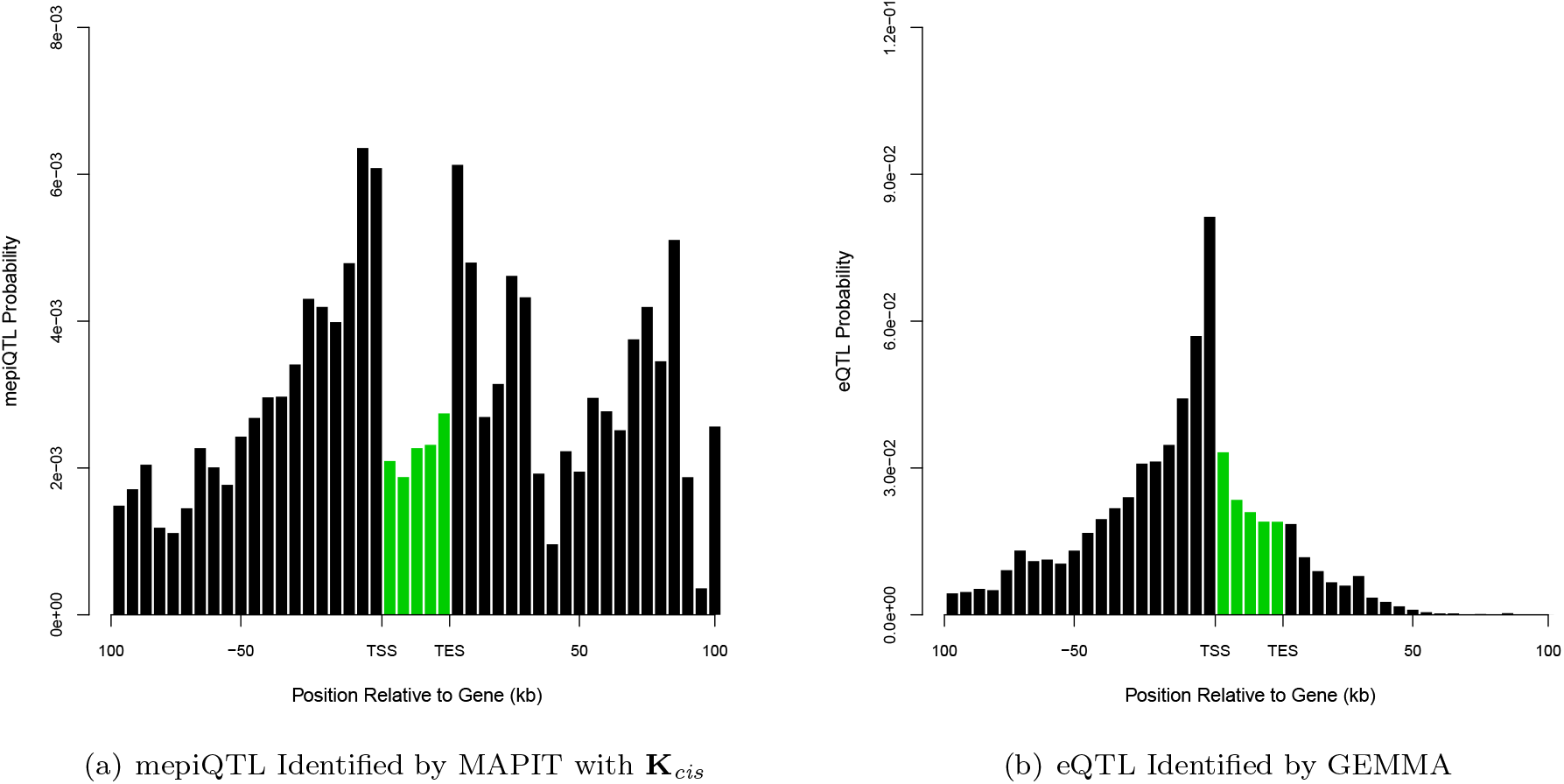
Enrichment of eQTL and mepiQTL SNPs in GEUVADIS data set. Shown here are the distribution of locations for significant SNPs, relative to the 5′ most gene transcription start site (TSS) and the 3‱ most gene transcription end site (TES). Figure (a) displays the marginally epistatic QTL (mepiQTL) detected by MAPIT using genetic relatedness matrix **K***_cis_*. Figure (b) corresponds to the expression QTL (eQTL) identified by the single-SNP via GEMMA. The x-axis of each plot divides a typical *cis*-candidate region into a series of bins. The y-axis plots the number of SNPs in each bin that have a p-value less than a gene specific Bonferroni-corrected significance p-value threshold *P* = 0.05/∑*_i_ s_i_*, where *s_i_* is the number of *cis*-SNPs for gene *i*, divided by the total number of SNPs in that bin. Bars in green denote the region bounded by the TSS and TES, with gene lengths divided into 20 bins for visibility — because the gene body is thus artificially enlarged, SNP density within genes cannot be directly compared with SNP density outside of genes.

Since MAPIT produces a single p-value for each tested variant, we may visualize the specific genomic locations that exhibit epistasis. Figure 7 displays zoomed-in manhattan plots for three of the twenty-two chromosomes where some of the most notable significant marginal epistatic effects were detected under the marginal genome-wide significance threshold (*P* = 1.828 × 10^−8^). Figures depicting the genome-wide epistatic scans of the other chromosomes can be found in Supporting Information (see S19 Fig.). Note that since there could be multiple p-values for a single SNP (i.e. its association evidence for multiple genes), we choose to display the minimum p-value as the summary statistic for any given SNP. We stress that the interpretation of these images is slightly different than what is used for traditional manhattan plots. Specifically, in these figures, spikes across chromosomes suggest loci where members involved in epistatic interactions can be found.

**Figure 7.**
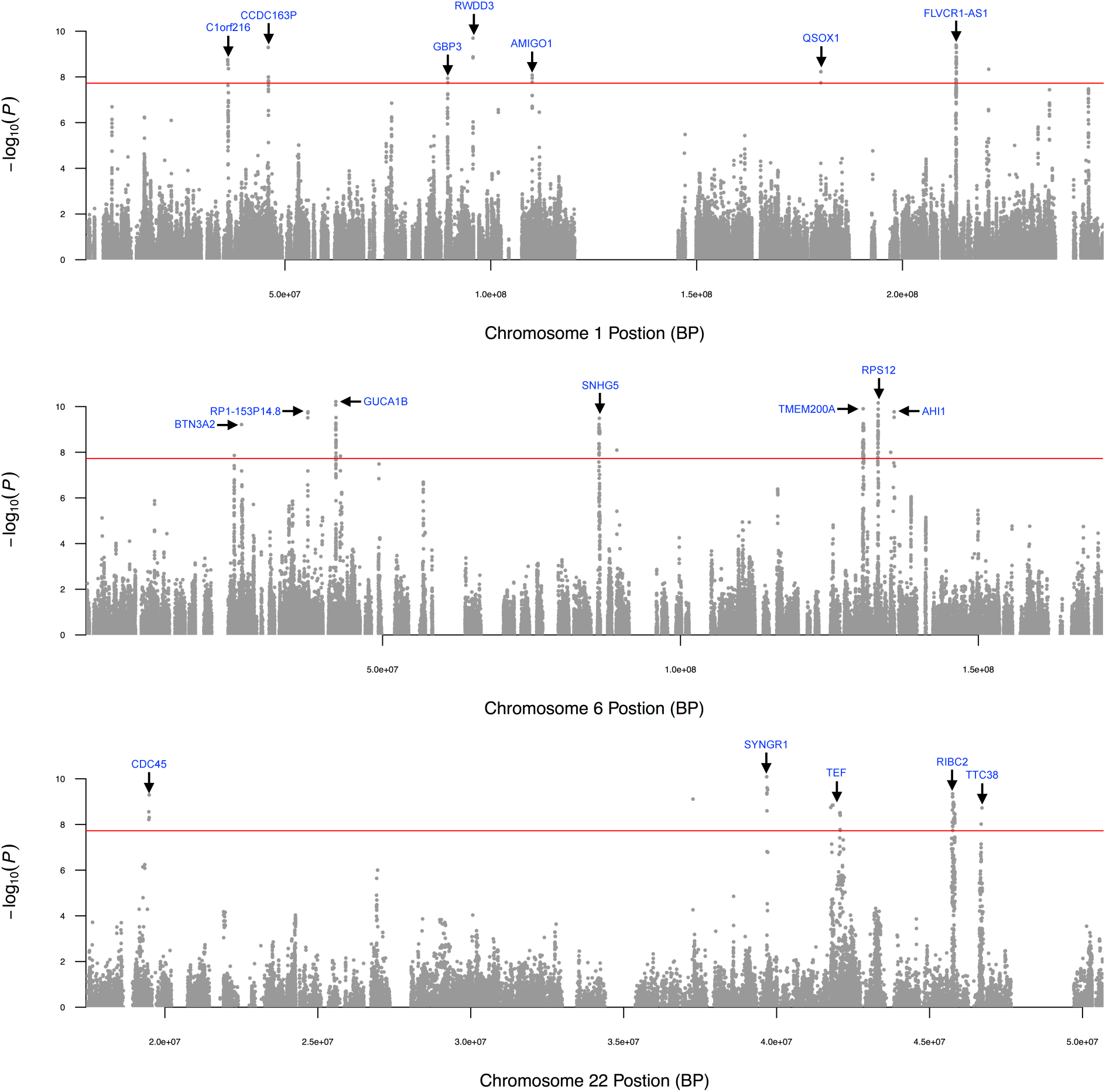
Select chromosome-wide scans for epistatic effects in GEUVADIS data set. Depicted are the −log_10_(*P*) transformed MAPIT p-values of quality-control-positive *cis*-SNPs plotted against their genomic position. Note that MAPIT was implemented with **K***_cis_*. Here, the epistatic associated genes are labeled (blue). The (red) horizontal line indicates a genome-wide significance threshold (*P* = 1.828 × 10^−8^). Note that all panels are truncated at log_10_(*P*) = 10 for consistency and presentation, although for some genes there are strongly marginally epistatic associated markers with values *P* ≈ 0.

In order to search for exact epistatic pairs among cis-SNPs through the filtering approaches, we took all significant mepiQTL and eQTL, and analyzed the pairwise interactions between them. This comparison allows us to assess the power of MAPIT and GEMMA as filtering approaches within the context of real data. Similar to what was done in the simulation studies, both filtering procedures were carried out by first ranking SNPs by either their mepiQTL or eQTL p-values, and then searching for associated epistatic pairs between the top *v* SNPs. Interactions between two mepiQTL or eQTL were then called significant if they had a joint p-value below a gene specific Bonferroni corrected threshold. Briefly, this threshold is computed as *P* = 0.05/(∑*_i_ q_i_*(*q_i_* − 1)/2), where *q_i_* is the number of top variants (among the top *v* SNPs) located in the *cis*-window of gene *i*. Once again, we applied the fully exhaustive search model in PLINK as a baseline power comparison, in which we examined all of the *cis*-SNP pairs for each gene. Note that for PLINK we only consider cis-SNP combinations as it was not computationally feasible to perform an absolute complete exhaustive search (i.e. every gene pair between all 1.2 million SNPs for each of the 16,000 genes). Figure 5(b) (and S17(d), S17(e), and S17(f) Fig.) depicts the number of significant pairwise interactions identified by MAPIT (with different **K** matrices) and GEMMA when searching between the top *v* = {1000, 2500, 5000, 7500, 10000, 15000, 20000} marginally associated variants. Altogether, the exhaustive search method in PLINK identified 7,361 significant epistatic pairs before the correction of residual confounding effects, and 8,643 significant pairwise interactions after the correction. These benchmarks are represented by the dotted lines in Figure 5(b), S17(d), S17(e), and S17(f), respectively.

As demonstrated in the numerical experiments, using MAPIT as the initial filtration procedure (as opposed to the single-SNP additive model) efficiently provides more power to finding significant epistatic pairs. Moreover, considering interactions amongst just the top 2,500 mepiQTL (in any of the considered settings for MAPIT) almost results in the same total number of significant epistatic pairs that is to be discovered by the fully exhaustive search model from PLINK. As previously shown, these results may be due to MAPIT's ability to marginally detect the “hub” SNPs of interactions. For example, under the expression for gene *C16orf88*, the SNP rs11645910 (MAPIT *P* ≈ 0) is a member of many of the top significant epistatic pairs. We also note that 36 of the 228 epistatic associated genes identified by MAPIT with **K***_cis_* have been verified in previous analyses on a different RNAseq data set from the TwinsUK cohort [37]. Similarly, 26 of the 120 mepiGenes discovered by MAPIT with **K***_trans_* and **K***_GW_*, and 35 of the 184 mepiGenes discovered by MAPIT with **K***_Pop_* after correction for residual confounders, have been verified in the same study.

In order to better explain why MAPIT was able to identify significant epistatic pairs in this study, we refer back to the variance component modeling approach [44, 83, 84] we used in simulations to evaluate the overall contribution of pairwise epistasis to the PVE for the expression of each gene (details in Methods and Material). Again, the basic idea behind dissecting the makeup of the PVE is using a linear mixed model with multiple variance components to partition the phenotypic variance into two distinct components: a linear component and a pairwise interaction component. Disregarding any random noise, we quantify the contribution of the two components by examining the proportion of PVE (pPVE) explained by that component. In Supporting information, we illustrate the estimates of the pPVE decomposition by additive effects and pairwise epistasis for all genes (see S20 Fig.). In this particular data set, we find that the mean pPVE for the pairwise interaction component is approximately 10%, which is consistent with previous studies on the same data [96]; while the additive effects only explain about 3.5% on average. To put this into better context, for the gene *C16orf88*, pairwise epistatic effects are estimated to explain approximately 5% of the PVE, while additive effects are estimated to only account for 3 × 10^−3^%. In fact, the pairwise order component actually explains a larger proportion of phenotypic variance than the linear component in the expression of 7,704 out of 15,607 genes.

Finally, we also want to point out one important caveat for mapping epistasis in real data: in the analyses of genetic mapping studies, apparent epistasis inferred by any epistatic mapping methods can sometimes be explained by same-locus additive effects [41]. This means that the results from all methods (MAPIT or the exhaustive search procedure or the additive-effect based filtering procedure) could be confounded by additive effects of untyped SNPs or uncontrolled SNPs in the same region, even though the power comparison among these methods remains fair. Dealing with these contingencies is difficult because it is impossible to precisely control for the additive fixed effects of all SNPs that reside in the same locus. Therefore, we caution against the over-interpretation of our analysis results in GEUVADIS and simply use the GEUVADIS data as an illustration on how MAPIT can be used as a first step towards understanding the genetic architecture of phenotypic variation.

## Discussion

We have presented MAPIT, a novel method and strategy for detecting variants that are involved in epistasis in genetic mapping studies. For each variant in turn, MAPIT estimates and tests its marginal epistatic effect — the combined epistatic effect between the examined variant and all other variants. By modeling and inferring the marginal epistatic effects, MAPIT can identify variants that exhibit non-zero epistatic interactions with any other variant without the need to identify the specific marker combinations that drive the epistatic association. Therefore, MAPIT represents an attractive alternative to standard methods [54–56, 63] for mapping epistasis. With both simulations and real data applications, we have illustrated the benefits of MAPIT.

In the present study, we have focused on estimating and testing marginal epistatic effects in the presence of pairwise interactions with MAPIT. MAPIT can also be easily extended to detect variants that are involved in higher-order interactions. Specifically, in the presence of higher-order interactions, we can introduce extra random effects terms to represent the combined higher-order interaction effects between the examined variant and all other variants — this would simply mean adding an extra random effects term for each extra higher-order of interactions. Under the normality assumption of the interaction effect sizes, the introduced random effects terms would all follow multivariate normal distributions, with the covariance matrices determined as a function of the Hadamard product of the additive genetic relatedness matrix [44, 84, 97]. Therefore, we can use a multiple variance component model with additional variance components to map epistatic variants in the presence of higher-order interactions. From there, we can test the variance components jointly to identify variants that are involved in any order of epistatic interactions. We can test each variance component separately to identify variants that are involved in a particular order epistatic interaction. Or, better still, we can perform variable selection on the variance components to identify which higher order interaction a particular variant of interest is involved in. Extending MAPIT to mapping high-order interactions will likely provide further insights into the epistatic genetic architecture of various traits and diseases.

We have focused on mapping epistasis for quantitative traits. For case-control studies, one may be tempted to follow previous approximate approaches of treating binary phenotypes as continuous traits and apply MAPIT directly [67, 98]. However, it would be desirable to extend MAPIT to accommodate case-control data or other discrete data types in a principled way. Two variance component models are available for handling case-control data, and each has its advantages and drawbacks. The first model is the liability threshold model, which models the liability score underlying the binary trait with a variance component model [79, 98]. The liability threshold model has been widely used for estimating heritability of common diseases, but relies on an asymptotically normal test, which, as is evident with our simulations, may fail to properly control for type I error at the genome-wide significance level for association tests. The second model is the logistic mixed model that has been well established in the statistics literature [99–102], and has been recently applied to perform association tests in case-control studies [103] as well as in RNA sequencing and bisulfite sequencing studies [104, 105]. However, unlike the liability threshold model, it is not straightforward to define and partition the phenotypic variance into various genetic components under the logistic mixed model. Therefore, future studies are needed to extend MAPIT to case-control studies by either unifying the two models or developing new methods that can perform rigorous hypothesis tests while enabling genetic partitioning of phenotypic variance.

In its current form, we have focused on demonstrating MAPIT with a variance component model. The variance component model in MAPIT effectively assumes that the interaction effect between the examined variant and every other variant follows a normal distribution. This normality assumption and the resulting variance component model have been widely used in many areas of genetics. For example, variance component models are used in rare variant tests to combine the additive effects of multiple rare variants to improve association mapping power [64, 65]. Similarly, variance component models are used to jointly model all genome-wide SNPs at once for estimating SNP heritability [67, 69]. Studies have already shown that variance component models produce unbiased estimates regardless of whether or not the underlying effect sizes follow a normal distribution, and are reasonably robust even when the model is severely misspecified [67, 69]. However, like any statistical model, the inference results of variance component models can be affected when the modeling assumptions are not fully satisfied. For example, recent studies have shown that the linkage disequilibrium (LD) pattern of causal SNPs can cause estimation bias that is either minor allele frequency (MAF) mediated or non-MAF-mediated [50,75]. Such LD-biases likely affect MAPIT in a similar fashion. Therefore, adapting the approaches taken in [50, 75] or developing a more realistic modeling assumption will likely improve the robustness of MAPIT even further. Fortunately, MAPIT can easily be extended to incorporate other effect size assumptions. Indeed, the main idea in MAPIT of mapping marginal epistatic effects is not restricted to the particular variance component model we examine here, nor is it restricted to the normality assumption of the interaction effect sizes. Therefore, we can incorporate sparsity-inducing priors for effect sizes if the number of interaction pairs is known to be small *a priori*. Alternatively, we can use the recently developed hybrid effect size prior that has been shown to work well under a variety of effect size distributions [67]. Different interaction effect size assumptions can be advantageous under different genetic architectures and incorporating them in different scenarios will likely improve the power of MAPIT further.

We have only compared MAPIT with two commonly used epistatic mapping methods that include an exhaustive search method and an additive effect prioritization method. There are many other novel methods that have been developed recently [53, 57, 60–62]. For example, a recently proposed partition retention method partitions individuals into different genotype groups that are defined based on the pairwise or high-order combinations of their genotypes (e.g. a total of 81 genotype groups for four SNPs that each have three genotype classes) [106]. More specifically, this method first computes the phenotypic variance across these genotype groups, and then examines each SNP in the combination by testing whether the SNP adds a significant contribution to the phenotypic variance across the groups. Despite its computational intensity, the partition retention method produces promising results. It would thus be important to compare MAPIT with the partition retention method, as well as others, in the future.

There are many other potential extensions of MAPIT. We have only focused on analyzing one phenotype at a time in this study. However, it has been extensively shown that modeling multiple phenotypes can often dramatically increase power [68, 74]. Therefore, it would be interesting to extend MAPIT to take advantage of phenotype correlations to identify pleiotropic epistatic effects. Modeling epistasis in the context of multiple phenotypes could be highly non-trivial, as we need to properly model the shared epistatic components between phenotypes, in addition to the shared additive effects between phenotypes. Modeling strategies based on the multivariate linear mixed model (mvLMM) [68, 74] could be helpful here.

MAPIT is not without its limitations. Perhaps the most noticeable limitation is that MAPIT cannot be used to directly identify the interaction pairs that drive individual variant association. In particular, after identifying a variant involved in epistasis, it is still unclear which variants it interacts with. Thus, despite being able to identify SNPs that are involved in epistasis, MAPIT is unable to directly identify detailed interaction pairs. However, we argue that being able to identify variants that are involved in epistasis is often an important first step towards identifying and understanding detailed epistatic associations. In addition, being able to identify SNPs involved in epistasis allows us to come up with an initial likely set of variants that are worth further exploration. Indeed, we advertise a two-step *ad hoc* epistasis association mapping procedure. First, we identify individual SNP associations with MAPIT. Then, we focus on the most significant associations from the first step to further test all of the pairwise interactions among them in order to identify specific epistatic interactions. Unlike the previous filtering strategies that are commonly used in epistatic mapping, our two-step procedure is unique in the sense that the SNP set identified in our first step contains SNPs that already display strong epistatic effects with other variants. Therefore, our two-step procedure outperforms alternative filtering strategies in simulations and real data applications. Nonetheless, we caution that the two-step procedure is still *ad hoc* in nature and could miss important epistatic associations. Therefore, exploring statistical approaches that can unify the two steps would be an interesting area for future research. Besides this main limitation, we also note that MAPIT can be computationally expensive. MAPIT requires fitting a variance component model for every SNP in turn, and fitting variance component models are known to be computationally challenging [66, 68]. In this study, we use the recently developed MQS method for variance component estimation and testing. Compared with the standard REML method, MQS is computationally efficient, allows for exact p-value computation based on the Davies method, and is statistically more efficient than the REML estimates when the variance component is small [77] — a property that is particularly relevant here considering the marginal epistatic effect size is often small. MQS allows us to apply MAPIT to moderately sized genetic mapping studies with thousands of samples and millions of variants, which is otherwise impossible using any other variance component estimation methods. Still, new algorithms are likely needed to scale MAPIT up to datasets that orders of magnitude larger in size.

## Acknowledgements

LC is supported by the National Science Foundation Graduate Research Program under Grant No. DGF-1106401. SM would like to acknowledge the support of grants NSF IIS-1546331, NSF DMS-1418261, NSF IIS-1320357, NSF DMS-1045153, and NSF DMS-1613261. XZ would like to acknowledge the support of NIH Grants R01HG009124, R01HL117626 (PI Abecasis), R21ES024834 (PI Pierce), R01HL133221 (PI Smith), and a grant from the Foundation for the National Institutes of Health through the Accelerating Medicines Partnership (BOEH15AMP, co-PIs Boehnke and Abecasis). We thank Kris C. Wood for helpful comments on a previous version of the manuscript. This study also makes use of data generated by the Wellcome Trust Case Control Consortium (WTCCC). A full list of the investigators who contributed to the generation of the data is available from www.wtccc.org.uk. Funding for the WTCCC project was provided by the Wellcome Trust under award 076113 and 085475. Any opinions, findings, and conclusions or recommendations expressed in this material are those of the author(s) and do not necessarily reflect the views of any of the funders or supporters.

